# The genome of the water strider *Gerris buenoi* reveals expansions of gene repertoires associated with adaptations to life on the water

**DOI:** 10.1101/242230

**Authors:** David Armisén, Rajendhran Rajakumar, Markus Friedrich, Joshua B Benoit, Hugh M. Robertson, Kristen A. Panfilio, Seung-Joon Ahn, Monica F. Poelchau, Hsu Chao, Huyen Dinh, HarshaVardhan Doddapaneni, Shannon Dugan, Richard A. Gibbs, Daniel S.T. Hughes, Yi Han, Sandra L. Lee, Shwetha C. Murali, Donna M. Muzny, Jiaxin Qu, Kim C. Worley, Monica Munoz-Torres, Ehab Abouheif, François Bonneton, Travis Chen, Li-Mei Chiang, Christopher P. Childers, Andrew Graham Cridge, Antonin Jean Johan Crumière, Amelie Decaras, Elise M. Didion, Elizabeth Duncan, Elena N. Elpidina, Marie-Julie Favé, Cédric Finet, Chris G.C. Jacobs, Alys Jarvela, Emily J. Jennings, Jeffery W. Jones, Maryna P. Lesoway, Mackenzie Lovegrove, Alexander Martynov, Brenda Oppert, Angelica Lillico-Ouachour, Arjuna Rajakumar, Peter Nagui Refki, Andrew J. Rosendale, Maria Emilia Santos, William Toubiana, Maurijn van der Zee, Iris M. Vargas Jentzsch, Aidamalia Vargas Lowman, Severine Viala, Stephen Richards, Abderrahman Khila

**Affiliations:** Institut de Génomique Fonctionnelle de Lyon, Université de Lyon, Université Claude Bernard Lyon 1, CNRS UMR 5242, Ecole Normale Supérieure de Lyon, 46, allée d’Italie, 69364 Lyon Cedex 07, France; Department of Molecular Genetics & Microbiology and UF Genetics Institute, University of Florida, 2033 Mowry Road, Gainesville, Florida 32610-3610, USA; Department of Biological Sciences, Wayne State University, Detroit, Michigan 48202, USA; Department of Biological Sciences, McMicken College of Arts and Sciences, University of Cincinnati, 318 College Drive, Cincinnati OH 45221-0006 USA; Department of Entomology, University of Illinois at Urbana-Champaign, Urbana, Illinois 61801, USA; Institute for Zoology: Developmental Biology, University of Cologne, Zülpicher Str. 47b, 50674 Cologne, Germany; School of Life Sciences, University of Warwick, Gibbet Hill Campus, Coventry CV4 7AL, UK; USDA-ARS Horticultural Crops Research Unit, 3420 NW Orchard Avenue, Corvallis, OR 97330, USA; Department of Crop and Soil Science, Oregon State University, 3050 SW Campus Way, Corvallis, OR 97331, USA; USDA Agricultural Research Service, National Agricultural Library, Beltsville, MD 20705, USA; Human Genome Sequencing Center, Department of Human and Molecular Genetics, Baylor College of Medicine, One Baylor Plaza, Houston, Texas 77030, USA; Howard Hughes Medical Institute, University of Washington, Seattle, WA 98195, USA; Lawrence Berkeley National Laboratory, Berkeley, California, USA; McGill University, Department of Biology, 1205 Avenue Docteur Penfield Avenue, Montréal, Québec H3A 1B1, Canada; Laboratory for Evolution and Development, Department of Biochemistry, University of Otago, P.O. Box 56, Dunedin, Aotearoa-New Zealand; School of Biology, Faculty of Biological Sciences, University of Leeds, Leeds LS2 9JT, United Kingdom; A.N. Belozersky Institute of Physico-Chemical Biology, Moscow State University, Moscow 119991, Russia; Institute of Biology, Leiden University, Sylviusweg 72, 2333 BE Leiden, Netherlands; Max Planck Institute for Chemical Ecology, Hans-Knöll Strasse 8, 07745 Jena, Germany; University of Maryland, College Park, 4291 Fieldhouse Dr., Plant Sciences Building, Rm. 4112, College Park, MD 20742, USA; Smithsonian Tropical Research Institute, Naos Island Laboratories, Apartado Postal 084-303092, Balboa, Ancon, Republic of Panama.; Department of Cell and Developmental Biology, University of Illinois, Urbana, IL.; Center for Data-Intensive Biomedicine and Biotechnology, Skolkovo Institute of Science and Technology, Skolkovo 143025 Russia; USDA Agricultural Research Service, Center for Grain and Animal Health Research, 1515 College Ave., Manhattan, KS, 66502 USA; Max-Planck-Institut für Evolutionsbiologie. Department of Evolutionary Genetics. August-Thienemann-Straße 2, 24306 Plön, Germany

## Abstract

The semi-aquatic bugs conquered water surfaces worldwide and occupy ponds, streams, lakes, mangroves, and even open oceans. As such, they inspired a range of scientific studies from ecology and evolution to developmental genetics and hydrodynamics of fluid locomotion. However, the lack of a representative water strider genome hinders thorough investigations of the mechanisms underlying the processes of adaptation and diversification in this group. Here we report the sequencing and manual annotation of the *Gerris buenoi (G. buenoi)* genome, the first water strider genome to be sequenced so far. *G. buenoi* genome is about 1 000Mb and the sequencing effort recovered 20 949 predicted protein-coding genes. Manual annotation uncovered a number of local (tandem and proximal) gene duplications and expansions of gene families known for their importance in a variety of processes associated with morphological and physiological adaptations to water surface lifestyle. These expansions affect key processes such as growth, vision, desiccation resistance, detoxification, olfaction and epigenetic components. Strikingly, the *G. buenoi* genome contains three Insulin Receptors, a unique case among metazoans, suggesting key changes in the rewiring and function of the insulin pathway. Other genomic changes include wavelength sensitivity shifts in opsin proteins likely in association with the requirements of vision in water habitats. Our findings suggest that local gene duplications might have had an important role during the evolution of water striders. These findings along with the *G. buenoi* genome open exciting research opportunities to understand adaptation and genome evolution of this unique hemimetabolous insect.

## Background

The semi-aquatic bugs (Gerromorpha) are a monophyletic group of predatory heteropteran insects characterized by their ability to live at the water-air interface ^1-4^. The Gerromorpha ancestor transitioned from terrestrial habitats to the water surface over 200 million years ago, and subsequently radiated into over 2 000 known species classified in eight families ^1^. The ancestral habitat of the Gerromorpha, as inferred from phylogenetic reconstruction, is humid terrestrial or marginal aquatic ^1,5,6^. Many lineages, such as water striders, became true water surface dwellers and colonized a diverse array of niches including streams, lakes, ponds, marshes, and even the open ocean ^1,7,8^. The invasion of this new habitat provided access to resources previously underutilized by insects and made the Gerromorpha the dominant group of insects at water surfaces. This novel specialized life style makes the Gerromorpha an exquisite model system to study how new ecological opportunities can drive adaptation and species diversification^2,9-11^.

The shift in habitat exposed these insects to new selective pressures that are divergent from their terrestrial ancestors. The Gerromorpha face two primary challenges unique among insects: how to remain afloat and how to generate efficient thrust on the fluid substrate ^2,3,12^. The bristles covering the legs of water striders, owing to their specific arrangement and density, act as a non-wetting structures capable of exploiting water surface tension by trapping air between the leg and water surface and keeping them afloat (Figure 1A) ^2,3,12,13^. Locomotion, on the other hand, is made possible through changes in the morphology and the patterns of leg movement (Figure 1B) ^2,3,12,13^ Two modes of locomotion are employed: an ancestral mode using the tripod gait through alternating leg movements, and a derived mode using the rowing gait through simultaneous sculling motion of the pair of middle legs (Figure 1B) ^2,12^. The derived mode through rowing is characteristic of water striders and is associated with a derived body plan where the middle legs are the longest (Figure 1A-B) ^2,12^. The specialization in water surface life is thought to be associated with new predator (Figure 1C) and prey (Figure 1D) interactions that shaped the evolutionary trajectory of the group. Other adaptations following invasion of water surfaces include their visual system to adapt to surface-underwater environment, wing polymorphism in relation with habitat quality and dispersal (Figure 1E) ^14^, and cuticle composition and its role in water exchange to counter water gain associated with living on water.

**Figure 1.**
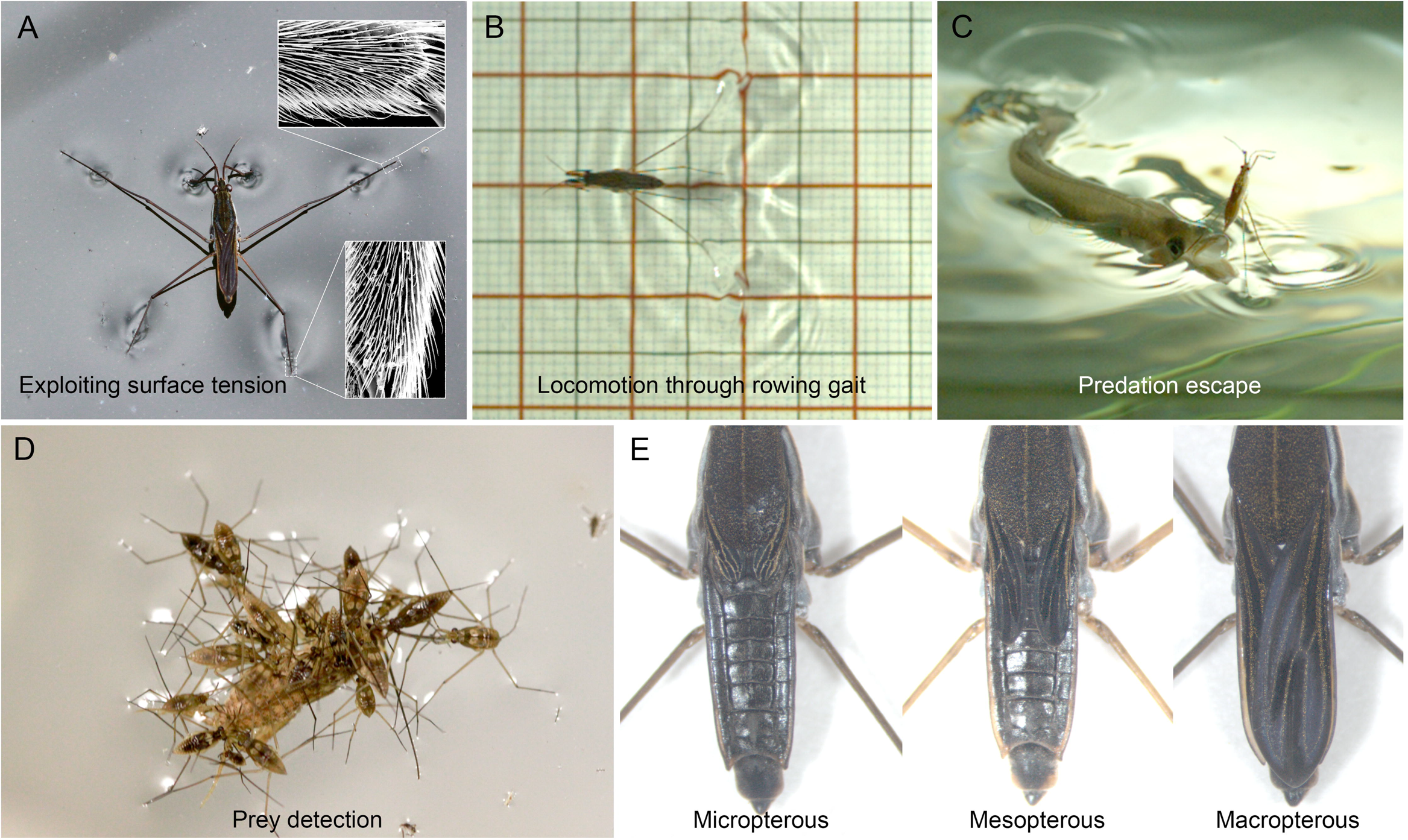
Aspects of the biology of water striders. **(A)** Adult *Gerris* on water and zoom in on the bristles allowing this adaptation using Scanning Electron Microscopy (insets). **(B)** *Gerris* rowing on the water surface, illustrating the adaptive locomotion mode. (C) Water strider jumping using its long legs to escape the strike of a surface hunting fish. (D) Hoarding behavior in water striders consisting of multiples individuals feeding on a cricket trapped by surface tension. (E) Wing polymorphism in *Gerris*, here illustrated by three distinct morphs with regard to wing size.

While we are starting to understand some developmental genetic and evolutionary processes underlying the adaptation of water striders to the requirements of water surface locomotion, prey-predator, and sexual interactions ^2,15-19^, studies of these mechanisms at the genomic level are hampered by the lack of a representative genome. Here we report the genome of the water strider *G. buenoi*, the first sequenced member of the Gerromorpha infra-order. *G. buenoi* is part of the Gerridae family, and has been previously used as a model to study sexual selection and developmental genetics ^15,20-22^. Moreover, *G. buenoi* can easily breed in laboratory conditions and is closely related to several other *G*. species used as models for the study of the hydrodynamics of water walking, salinity tolerance, and sexual conflict. With a particular focus on manual annotation and analyses of processes involved in phenotypic adaptations to life on water, our analysis of the *G. buenoi* genome hints that the genomic basis of water surface invasion might be, at least in part, linked to local gene duplications.

## Results and discussion

### General features of the *G. buenoi* genome

The draft assembly of *G. buenoi* genome comprises 1 000 194 699 bp (GC content: 32.46%) in 20 268 scaffolds and 304 909 contigs (N50 length 344 118 bp and 3 812 bp respectively). The assembly recovers ∼87 % of the genome size estimated at ∼1.15 GB based on kmer analysis. *G.buenoi* genome is organized in 18 autosomal chromosomes with a XX/X0 sex determination system ^23^. MAKER automatic annotation pipeline predicted 20 949 protein-coding genes; a number that is higher than the 16 398 isogroups previously annotated in the transcriptome of the closely related species *Limnoporus dissortis* (PRJNA289202) ^18,24^, the 14 220 genes in bed bug *Cimex lectularius* genome ^25^ and the 19 616 genes in milkweed bug *Oncopeltus fasciatus* genome ^26^. The final *G. buenoi* OGS 1.0 includes 1 286 manually annotated genes representing development, growth, immunity, cuticle formation as well as olfaction and detoxification pathways (see Supplementary Material). Using OrthoDB (http://www.orthodb.org) ^27^, we found that 77.24% of the *G. buenoi* genes have at least one orthologue in other arthropod species (Figure **2**). We then used benchmarking sets of universal single-copy orthologs (BUSCOs) ^28^ to assess the completeness of the assembly. A third of BUSCOs (31%) were missing and 28.6% were fragmented, which correlates with the high number of gaps observed in the draft assembly (Supplementary Tables 1 and 2). On the other hand, 2.2 % of BUSCOs showed signs of duplication but functional GO term analysis showed no particular function enrichment.

**Figure 2.**
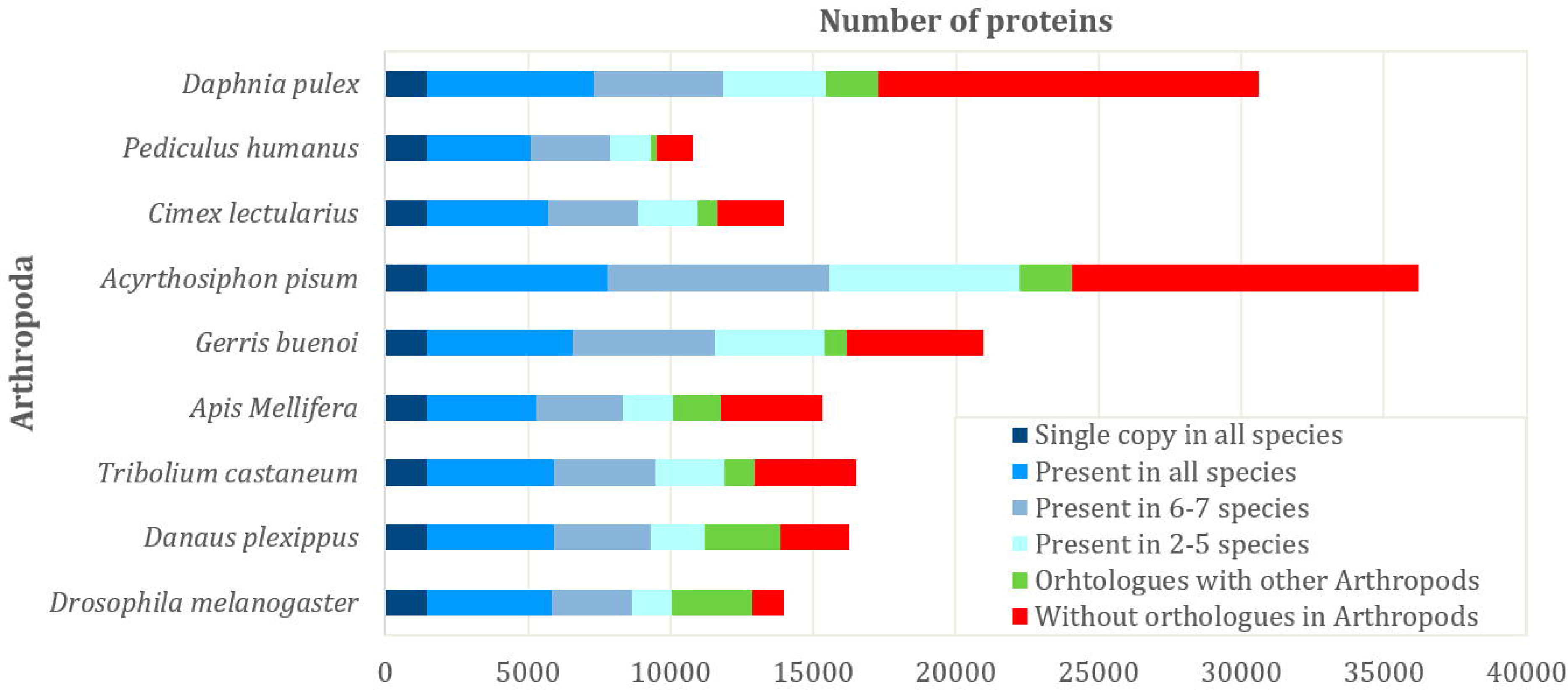
Orthology comparison between *Gerris buenoi* and other arthropod species. Genome proteins were clustered with proteins of other 12 arthropod species based on OrthoDB orthology.

In addition to BUSCOs, we used Hox and Iroquois Complex (Iro-C) gene clusters as indicators of draft genome quality and as an opportunity to assess synteny among species. The Hox cluster is conserved across the Bilateria ^29^, and the Iro-C is found throughout the Insecta ^25,30^. In *G. buenoi*, we were able to find and annotate gene models for all ten Hox genes (Supplementary Table 3). While linkage of the highly conserved central class genes *Sex combs reduced, fushi tarazu*, and *Antennapedia* occurred in the expected order and with the expected transcriptional orientation, the linked models of *proboscipedia* and *zerknüllt (zen)* occur in opposite transcriptional orientations (head-to-head, rather than both 3’ to 5’). Inversion of the divergent *zen* locus is not new in the Insecta ^31^, but was not observed in the hemipteran *C. lectularius*, in which the complete Hox cluster was fully assembled ^25^. Future genomic data will help to determine whether such microinversion within the Hox cluster is conserved within the hemipteran family Gerridae. Assembly limitations are also manifest in that the complete gene model for *labial* is present but split across scaffolds, while only partial gene models could be created for *Ultrabithorax* and *Abdominal-B*. For the small Iroquois complex, clear single copy orthologues of both *iroquois* and *mirror* are complete but not linked in the current assembly (Supplementary Table 3). However, both genes are located near the ends of their scaffolds, and direct concatenation of the scaffolds (5’-Scaffold451-3’, 3’-Scaffold2206-5’) would correctly reconstruct this cluster: (1) with both genes in the 5’-to-3’ transcriptional orientation along the (+) DNA strand, (2) with no predicted intervening genes within the cluster, and (3) with a total cluster size of 308 Kb, which is fairly comparable with that of other recently sequenced hemipterans in which the Iro-C cluster linkage was recovered (391 Kb in *C. lectularius* ^25^ and 403 Kb in *O. fasciatus^26^*). Lastly, we examined genes associated with autophagy processes, which are highly conserved among insects, and all required genes are present within the genome (Supplementary Table 3). Therefore, Hox and Iroquois Complex (Iro-C) gene cluster analyses along with the presence of a complete set of required autophagy genes suggest a good gene representation and supports further analysis.

### Adaptation to water surface locomotion

One of the most important morphological adaptations that enabled water striders to conquer water surfaces is the change in shape, density, and arrangement of the bristles that cover the contact surface between their legs and the fluid substrate. These bristles, by trapping the air, act as a non-wetting structure that cushion between the legs and the water surface (Figure 1A)^2,3,12,13^. QTL studies in flies uncovered dozens of candidate genes and regions linked to variation in bristle density and morphology ^32. In the^ *G. buenoi* genome we were able to annotate 90 out of 120 genes known to be involved in bristle development ^32,33^ (Supplementary Table 4). Among those genes we found a single duplication, the gene *Beadex (Bx)*,

Similar duplication found in *C. lectularius* and *H. halys* suggest that *Bx* duplication occurred at their last common ancestor prior to Gerromorpha speciation. In *Drosophila, Bx* is involved in neural development by controlling the activation of *achaete-scute* complex genes ^34^ and mutants of *Bx* have extra sensory organs ^34^. We think it is reasonable to speculate that *Beadex* duplication might then have been exploited water striders and be linked to the changes in bristle pattern and density, thus opening new research avenues to further understand the adaptation of water striders to water surface life style.

### A unique addition to the Insulin Receptor family in Gerromorpha

The Insulin signalling pathway coordinates hormonal and nutritional signals in animals ^35-37^. This facilitates the complex regulation of several fundamental molecular and cellular processes including transcription, translation, cell stress, autophagy, and physiological states, such as aging and starvation ^37-40^. The action of Insulin signalling is mediated through the Insulin Receptor (InR), a transmembrane receptor of the Tyrosine Kinase class ^41^. While vertebrates possess one copy of the InR ^42^, arthropods generally possess either one or two copies. Interestingly, the *G. buenoi* genome contains three distinct InR copies, making it a unique case among metazoans. Further sequence examination using in-house transcriptome databases of multiple Gerromorpha species confirmed that this additional copy is common to all of them indicating that it has evolved in the common ancestor of the group (Figure 3). In addition to their presence in the transcriptomes of multiple species, cloning of the three InR sequences using PCR, indicates that these sequences originate from three distinct coding genes that are actively transcribed in this group of insects. Comparative protein sequence analysis revealed that all three InR copies possess all the characteristic domains found in the InR in both vertebrates and invertebrates (Figure 3A). To determine which of these three InR copies is the new addition to the *G. buenoi* genome, we performed a reconstruction of phylogenetic relationships between these sequences in a sample of eight Gerromorpha (three InR copies) and fifteen Insecta (one or two InR copies). This analysis clustered two InR copies into InR1 and InR2 distinct clusters (Figure 3B). Furthermore, Gerromorphan InR1 and InR2 copies clustered with bed bug and milkweed bug InR1 and InR2 respectively, while the Gerromorpha-restricted copy clustered alone (Figure 3B, Supplementary Figure 1). These data suggest that the new InR copy, we called InR1-like, originates from the InR1 gene much probably at time of Gerromorpha speciation. A closer examination of the organization of the genomic locus of InR1-like gene in *G. buenoi* genome revealed that this copy is intronless. This observation, together with the phylogenetic reconstruction, suggests that InR1-like is a retrocopy of InR1 that may have originated through RNA-based duplication ^43^.

**Figure 3.**
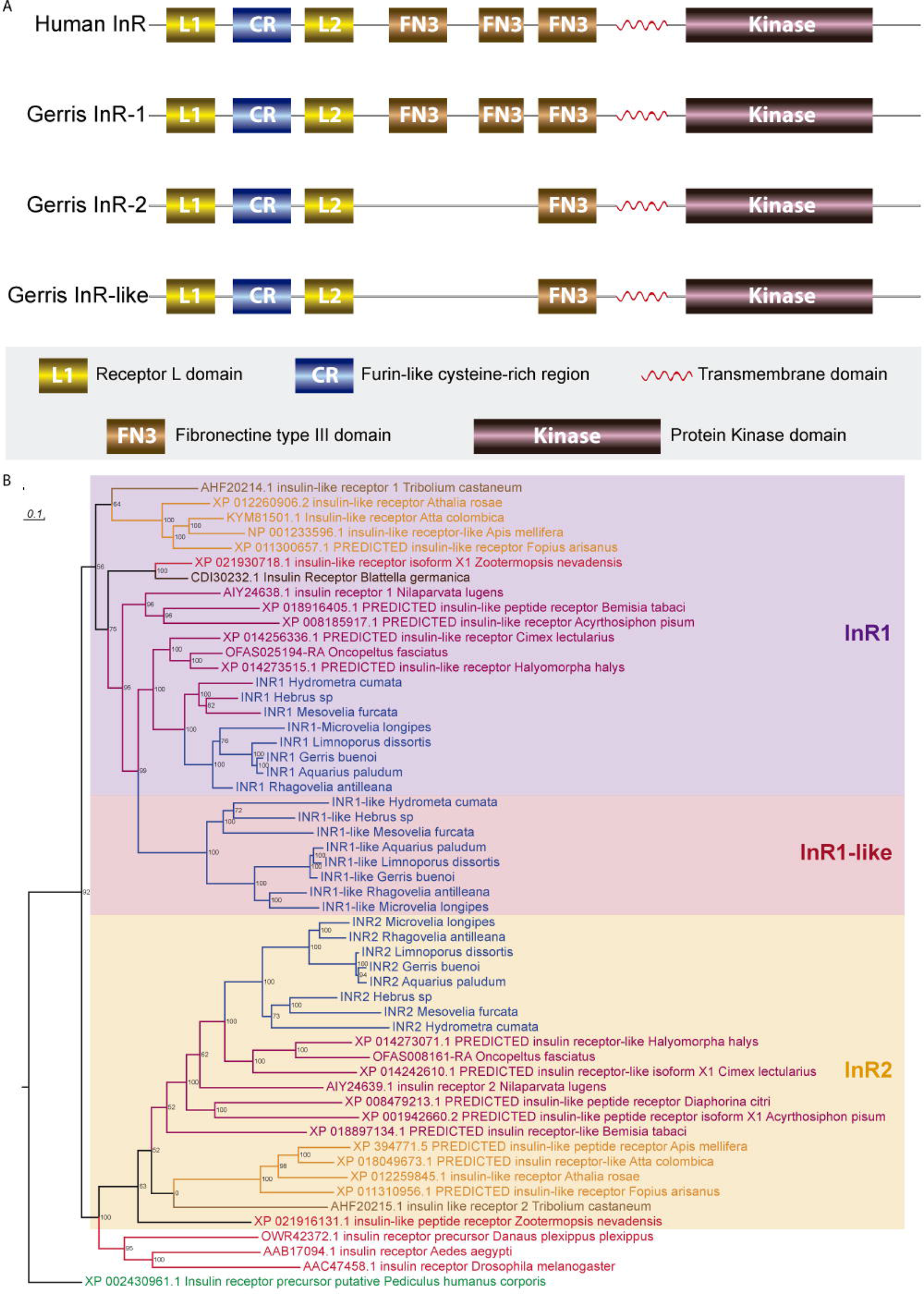
Characterization of the three copies of the Insulin Receptor in *Gerris buenoi*. **(A)** Protein domain comparison between the three InRs of *G. buenoi* and the Human InR. **(B)** InR phylogenetic relationship amongst Insecta. Color code: Blattodea (dark brown), Coleoptera (light brown), Diptera (red), Gerromorpha (blue), Hemiptera (except Gerromorpha)(purple), Hymenoptera (orange) and Phthiraptera (green). Branch support numbers at branches.

In insects, the Insulin signalling pathway has been implicated in the developmental regulation of complex nutrient-dependent growth phenotypes such as beetle horns and wing polyphenism in treehooppers, as well as morphological caste differentiation in social termites and bees ^44-47^. In the particular case of wing polymorphism in *G. buenoi* ^1,14,47^, our analysis found no DNA methylation signature as previously found in wing polyphenic ants and aphids ^48-52^ but an increased number of histone clusters and a unique histone methyltransferase *grappa* duplication (see Supplementary Data). It will thus be interesting to test the functional significance of the new InR copy in phenotypic plasticity of wing polyphenism but also how it impacts the role of the Insulin signalling in other aspects of *G. buenoi* biology and its involvement in the growth of other appendages.

### A lineage-specific expansion and possible sensitivity shifts of long wavelength sensitive opsins

The visual ecology and exceptionally specialized visual system of water striders has drawn considerable interest ^53,54^. Consisting of over 900 ommatidia, the prominent compound eyes of water striders are involved in prey localization, mating partner pursuit, predator evasion and, very likely, dispersal by flight ^55-57^. Realization of the first three tasks in the water surface to air interphase are associated with differences in the photoreceptor organization of the dorsal vs ventral eye ^53^, a lateral acute zone coupled to neural superposition ^58,59^ and polarized light-sensitive ^60^ (see Supplementary Data). Each water strider ommatidium contains 6 outer and 2 inner photoreceptors. Recent work has produced evidence of at least 2 types of ommatidia with either green (∼530nm) or blue (∼470-490nm) sensitive outer photoreceptors ^61^, but the wavelength specificity of the two inner photoreceptors cells is still unknown. At the molecular level, the wavelength-specificity of photoreceptor subtypes is in mostly determined by the expression of paralogous light sensitive G-protein coupled receptor proteins, opsins, that differ in their wavelength absorption maxima. Interestingly, our genomic analysis of opsin diversity in *G. buenoi* uncovered 8 opsin homologs. This included one member each of the 3 deeply conserved arthropod non-retinal opsin subfamilies (c-opsin, Arthropsin, and Rh7 opsin (see Supplementary Data)) and 5 retinal opsins (Figure 4A and Supplementary Figure 2). The latter sorted into one member of the UV-sensitive opsin subfamily and 4 tightly tandem clustered members of the long wavelength sensitive (LWS) opsin subfamily (Figure 4A). Surprisingly, both genomic and transcriptome search in *G. buenoi* and other water strider species failed to detect sequence evidence of homologs of the otherwise deeply conserved blue-sensitive opsin subfamily (Figure 4B; Supplementary Table 5) ^62^. Although the apparent lack of blue opsin in *G. buenoi* was unexpected given the presence of blue sensitive photoreceptors ^61^, it was consistent with the lack of blue opsin sequence evidence in available genomes and transcriptomes of other heteropteran species including *Halyomorpha halys, Oncopeltus fasciatus, Cimex lectularius, Rhodnius prolixus*. Blue opsin, however, is present in other hemipteran clades, including Cicadomorpha *(Nephotettix cincticeps)* and Sternorrhyncha *(Pachypsylla venusta)* (Figure 4B). Taken together, these data lead to the conclusion that the blue-sensitive opsin subfamily was lost early in the last common ancestor of the Heteroptera (Figure 4B and Supplementary Table 5). This raised the question of which compensatory events explain the presence of blue sensitive photoreceptors in water striders.

**Figure 4.**
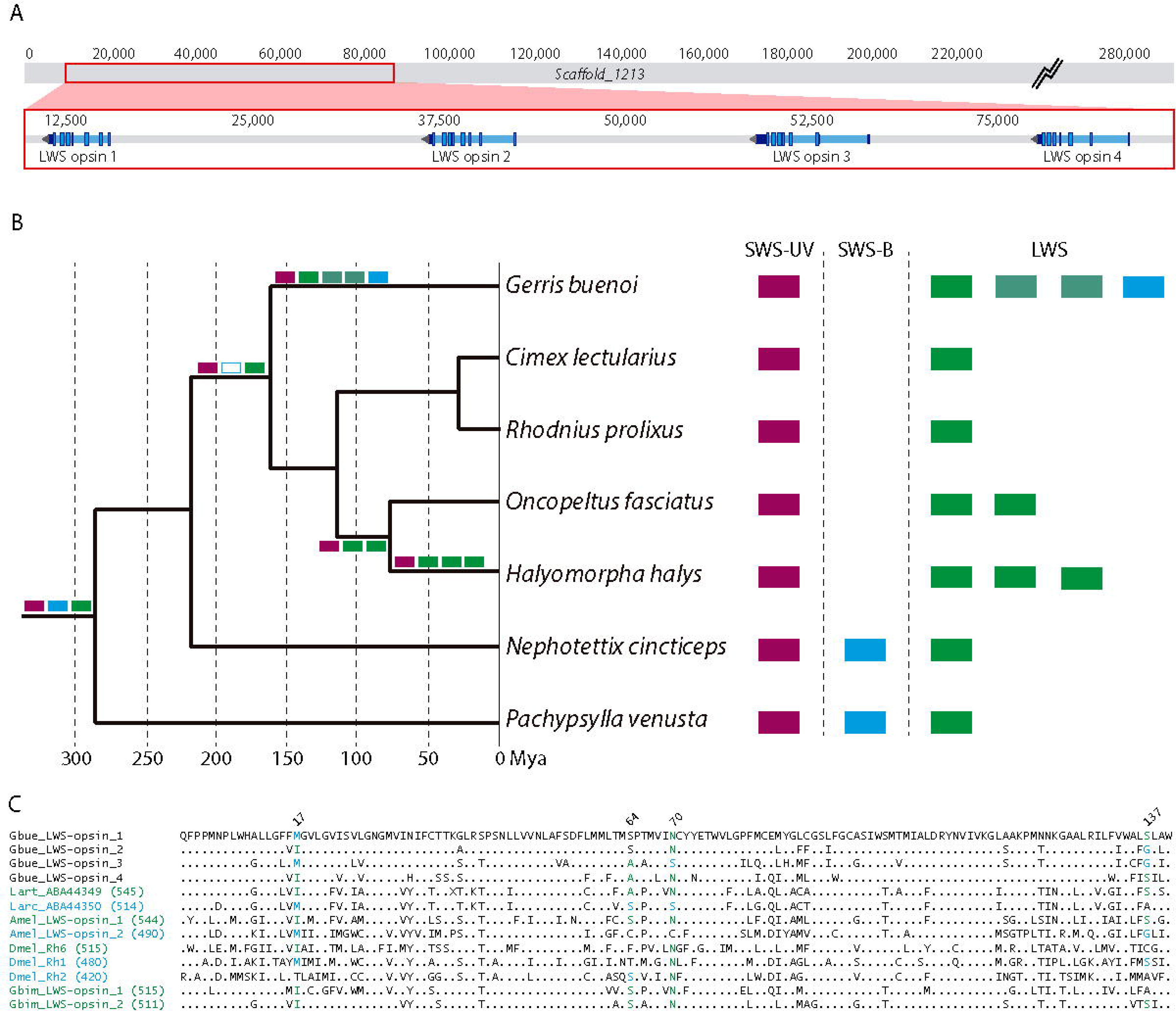
Genomic locus and global analysis of the *Gerris buenoi* opsin gene repertoire. **(A)** Structure of the scaffold containing the four *G.buenoi* long wavelength (LWS) opsins. **(B)** Retinal opsin repertoires of key hemipteran species and reconstructed opsin subfamily loss and expansion events along the hemipteran phylogeny. **(C)** Comparison of amino acid residues at the four tuning sites identified in the LWS opsins of Lepidoptera ^63,64^. Site numbers based on ^63^. Numbers in parentheses are experimentally determined sensitivity maxima. Species abbreviations: Amel = *Apis mellifera*, Dmel = *Drosophila melanogaster*, Gbue = *Gerris buenoi*, Gbim = *Gryllus bimaculatus*, Larc = *Limenitis archippus*, Lart = *Limenitis arthemis astyanax*.

Studies in butterflies and beetles produced evidence of blue sensitivity shifts in both UV- and LWS-opsin homologs following gene duplication ^63-65^. In butterflies, molecular evolutionary studies have implicated amino acid residue differences at four protein sequence sites in sensitivity shifts from green to blue: Ile17Met, Ala64Ser, Asn70Ser, and Ser137Ala ^63,64^ (Figure 4C, Supplementary Figure 2 and Supplementary Data). Based on sequence site information from physiologically characterized LWS opsins in other insect orders and the degree of amino acid residue conservation at these sites in a sample of 114 LWS opsin homologs from 54 species representing 12 insect orders (Supplementary Data File 1 and Supplementary Data) we could identify *G. buenoi* LWS opsin 3 as a candidate blue-shifted paralog with the highest confidence followed by *G. buenoi* LWS opsin 1 and 2. Moreover, the *G. buenoi* LWS opsin 4 paralog matches all of the butterfly green-sensitive amino acid residue states, thus favoring this paralog as green-sensitive (Figure 4). These conclusions are further backed by the fact that water striders lack ocelli, which implies that all four paralogs are expressed in photoreceptors of the compound eye. Overall, it thus seems most likely that the differential expression of the highly sequence-diverged *G. buenoi* LWS opsin paralogs accounts for the presence of both blue- and green-sensitive peripheral photoreceptors in water striders. Moreover, given that the outer blue photoreceptors have been specifically implicated in the detection of contrast differences in water striders ^61^, it is tempting to speculate that the deployment of blue-shifted LWS opsins represents another parallel to the equally fast-tracking visual system of higher Diptera in addition to open rhabdomeres, neural superposition, and polarized light-sensitivity.

### Expansion of cuticle gene repertoires

Desiccation resistance is essential to the colonization of terrestrial habitats by arthropods ^66^. However, contrary to most insects, the Gerromorpha spend their entire life cycle in contact with water and exhibit poor desiccation resistance ^1^. Cuticle proteins and aquaporins are essential for desiccation resistance through regulation of water loss and rehydration ^67-70^. In the *G. buenoi* genome, most members of cuticular and aquaporin protein families are present in similar numbers compared to other hemipterans (Supplementary Table 6 and Supplementary Figure 3). We identified 155 putative cuticle proteins belonging to five cuticular families: CPR (identified by Rebers and Riddiford Consensus region), CPAP1 and CPAP3 (Cuticular Proteins of Low-Complexity with Alanine residues), CPF (identified by a conserved region of about 44 amino acids), and TWDL (Tweedle) ^71,72^ (Supplementary Table 6). Interestingly, almost half of them are arranged in clusters suggesting local duplication events (Supplementary Table 7). Moreover, while most insect orders, including other hemipterans, have only three TWDL genes, we found that the TWDL family in *G. buenoi* has been expanded to 10 genes (Supplementary Figure 4). This expansion of the TWDL family is similar to that observed in some Diptera which contain *Drosophila*-specific and mosquito-specific TWDL expansions ^72,73^ and which *TwdlD* mutation alter body shape in *Drosophila* ^73^. Therefore, a functional analysis of TWDL genes and comparative analysis with other hemipterans will provide important insights into the evolutionary origins and functional significance of TWDL expansion in *G. buenoi*.

### Prey detection in water surface environments

Unlike many closely related species that feed on plants sap or animal blood, *G. buenoi* feeds on various arthropods trapped by surface tension (Figure 1D), thus making their diet highly variable. Chemoreceptors play a crucial role for prey detection and selection in addition to vibrational and visual signals. We annotated the three families of chemoreceptors that mediate most of the sensitivity and specificity of chemoperception in insects: Odorant Receptors (ORs; Supplementary Figure 5A), Gustatory Receptors (GRs; Supplementary Figure 5B) and Ionotropic Receptors (IRs; Supplementary Figure 5C) (e.g. ^74,75^). Interestingly, we found that the number of chemosensory genes in *G. buenoi* is relatively elevated (Supplementary Table 8). First, the OR family is expanded, with a total of 155 OR proteins. This expansion is the result of lineage-specific “blooms” of particular gene subfamilies, including expansions of 4, 8, 9, 13, 13, 16, 18, and 44 proteins, in addition to a few divergent ORs and the highly conserved OrCo protein (Supplementary Figure 5A and Supplementary Data). Second, the GR family is also fairly large (Supplementary Figure 5B), but the expansions here are primarily the result of large alternatively spliced genes, such that 60 genes encode 135 GR proteins (Supplementary Table 8). These GRs include 6 genes encoding proteins related to the carbon dioxide receptors of flies, three related to sugar receptors, and one related to the fructose receptor (Supplementary Figure 5B). The remaining GRs include several highly divergent proteins, as well as four blooms, the largest of which is 80 proteins (Supplementary Figure 5B and Supplementary Data). By analogy with *D. melanogaster*, most of these proteins are likely to be “bitter” receptors, although some might be involved in perception of cuticular hydrocarbons and other molecules. Finally, IR family is expanded to 45 proteins. In contrast with the OR/GR families where the only simple orthologs across these four heteropterans and *Drosophila* are the single OrCo and fructose receptor, the IR family has single orthologs in each species not only of the highly conserved co-receptors (IR8a, 25a, and 76b) but also receptors implicated in sensing amino acids, temperature, and humidity (Ir21a, 40a, 68a, and 93a). As is common in other insects the amine-sensing IR41a lineage is somewhat expanded, here to four genes, while the acid-sensing IR75 lineage is unusually highly expanded to 24 genes, and like the other heteropterans there are nine more highly divergent IRs (Supplementary Figure 5C and Supplementary Data).

We hypothesize that the high number of ORs may be linked to prey detection based on odor molecules in the air-water interface, although functional analysis will be needed to test the validity of this hypothesis. These receptors might help to complement water surface vibrations as prey detection system by expanding the spectrum of prey detectability. Similarly, being more scavengers than active hunters, *G. buenoi* are frequently faced with dead prey fallen on water for which they have to evaluate the palatability. As toxic molecules are often perceived as bitter molecules, the GR expansion might provide a complex bitter taste system to detect and even discriminate between molecules of different toxicities ^76^. Finally, expansion of the IR family could be linked with prey detection as well as pheromone detection of distant partners as IRs recognize, preferentially water-soluble hydrophilic acids and amines, many of which are common chemosensory signals for aquatic species^77,78^.

### Detoxification pathways

Water striders can be exposed to toxic compounds found in water, due to human activities, or in their prey either naturally or by exposure to pesticides or insecticides. Insect cytochrome P450 (CYP) proteins play a role in metabolic detoxification of xenobiotics including insecticides ^79,80^. They are also known to be responsible for the synthesis and degradation of endogenous molecules, such as ecdysteroids ^81^ and juvenile hormone ^82^. The insect CYPs comprise of one of the oldest and largest gene families in insects, which underwent great diversity from consecutive gene duplications and the subsequent diversification that extended the organism’s adaptive range ^83^. In addition to CYP proteins we have also surveyed the presence of UDP-glycosyltransferases (UGTs) genes in *G. buenoi*. UGTs are important for xenobiotic detoxification and the regulation of endobiotics in insects ^84^. UGTs catalyse the conjugation of a range of small hydrophobic compounds to produce water-soluble glycosides that can be easily excreted outside the body in a number of insects ^85,86^.

A total of 103 CYP genes (Supplementary Table 9 and Supplementary Data File 2) and 28 putative UGT genes including several partial sequences due to genomic gaps (Supplementary Table 10) were annotated and analyzed in the *G. buenoi* genome. Ten more CYP fragments were found, but they were not included in this analysis due to their short lengths (<250 aa). This is the largest number of CYP genes among the hemipteran species of which CYPomes were genome-widely annotated: *O. fasciatus* (58 CYPs), *R. prolixus* (88 CYPs) and *N. lugens* (68 CYPs) ^26,87,88^, as well as *D. melanogaster* (85), the *A. mellifera* (45), and *B. mori* (86) (Supplementary Table 9). Indeed, *G. buenoi* CYPs number is only exceeded by that of *T. castaneum* (131). CYP genes fall into one of the four distinct groups of CYP gene family, named the Clan 2, Clan mito, Clan 3 and Clan 4, where 6 genes 62 genes 25 genes, and 10 genes are present, respectively (Figure 5) (see Supplementay Data). Similarly, the number of UGT genes is also higher than that of *O. fasciatus* (1) ^26^, *C. lectularius* (7) ^25^, and *D. melanogaster* (11), *A. mellifera* (6) and *B. mori* (14) ^89^ and identical to *T. castaneum* (28) ^89^.

**Figure 5.**
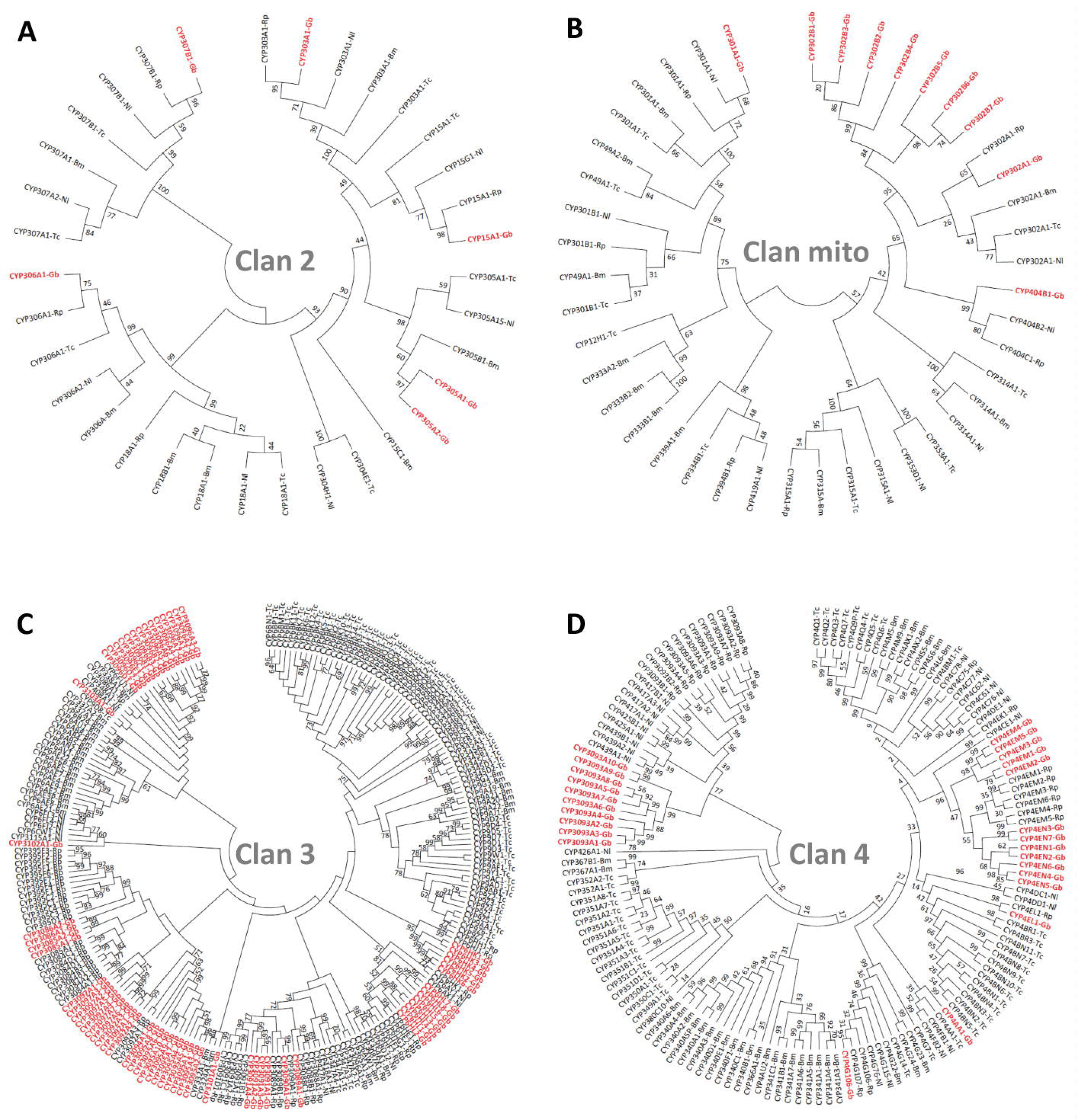
Phylogenetic analysis of four different Clans of the cytochrome P450s of *Gerris buenoi* with other insect species. (A) Clan 2, (B) Clan mitochondria, (C) Clan 3, and (D) Clan 4. The *G. buenoi* sequences are indicated in red and bold.

Interestingly, in both cases CYP and UGT gene family expansions seems to closely linked with tandem duplication events. In the particular case of *G. buenoi* CYPs, the Clan 2 and Clan mito have undergone relatively little gene expansion, probably conserved for essential roles in endogenous and/or exogenous metabolism (Figure 5A and B). However, an exceptional gene expansion is observed in the mitochondrial Clan of the *G. buenoi* CYPs, where seven CYP302Bs form a lineage-specific cluster (Figure 5B). The Clan 3 and Clan 4 are highly expanded in insect CYPs, as observed in insects such as *T. castaneum, B. mori, R. prolixus*, and *N. lugens* and show a high degree of gene expansion in *G. buenoi*, of which 45% (28/62 CYP genes) might have been generated by tandem gene duplications (Figure 5C and D). On the other hand, 10 UGT genes are arrayed in a row in Scaffold1549 suggesting gene duplication events may have produced such a large gene cluster (Supplementary Figure 6). In addition, multiple genes lie in Scaffold1323, Scaffold3228, and Scaffold2126 with 4, 3, and 2 UGT genes, respectively. A consensus Maximum-likelihood tree (Supplementary Figure 7) constructed with conserved C-terminal half of the deduced amino acid sequences from *G. buenoi* UGTs supports the conclusion that clustered genes placed in the same genomic location are produced by gene duplication.

Overall, phylogenetic analysis revealed not only CYPs and UGTs conserved orthology in insects, but also their lineage-specific gene expansions probably via lineage-specific gene duplication. We hypothesize that such expansion has been important to diversify xenobiotic detoxification range and the regulation of endobiotics during the transition to water surface niches.

## Conclusions

The sequencing of *G. buenoi* genome provides a unique opportunity to understand the molecular mechanisms underlying adaptations to water surface life and the diversification that followed. In particular, gene duplication is known to drive the evolution of adaptations and evolutionary innovations in a variety of lineages including water striders ^90-93^. The *G. buenoi* genome revealed a number of local and cluster duplications in genes that can be linked to processes associated with the particular life style of water striders. Some are shared with close related Hemiptera like for example, *Beadex*, an activator of *Achaete/Scute* complex known to play an important role in bristle development, present in two copies in the *G. buenoi* genome. Other genes and gene families duplications are unique, like Insulin Receptors involved in a range of processes including wing development, growth and scaling relationships and a number of life history traits such as reproduction ^44,47,94^. Functional significance of histone methyltransferase *grappa* and histone clusters duplications remains unknown opening new avenues in the study of the relationship between epigenetics and phenotypic plasticity. Expansions in the cuticle protein families involved in desiccation resistance or genes repertoires involved in xenobiotic detoxification and endobiotic regulation pathways may have had an important role during the specialization in water surface habitats ^73,95^. The expansion of the opsin gene family and possible sensitivity shifts are also likely associated with particularities of polarized light sensitivity due to the water environment where *G. buenoi* specializes. The impact of these duplications on the adaptation of water striders to water surface habitats remains to be experimentally tested. *G. buenoi*, as a recently established experimental model, offers a range of experimental tools to test these hypotheses. The *G. buenoi* genome, therefore, provides a good opportunity to begin to understand how lineages can burst into diversification upon the conquest of new ecological habitats.

## Methods

### Animal collection and rearing

Adult *G. buenoi* individuals were collected from a pond in Toronto, Ontario, Canada. *G. buenoi* were kept in aquaria at 25 °C with a 14-h light/10-h dark cycle, and fed on live crickets. Pieces of floating Styrofoam were regularly supplied to female water striders to lay eggs. The colony was inbred following a sib-sib mating protocol for six generations prior to DNA/RNA extraction.

### DNA and total RNA extraction

Genomic DNA was isolated from adults using Qiagen Genome Tip 20 (Qiagen Inc, Valencia CA). The 180bp and 500bp paired-end libraries as well as the 3kb mate-pair library were made from 8 adult males. The 8kb mate-pair library was made from 6 adult females. Total RNA was isolated from 39 embryos, three first instar nymphs, one second instar nymph, one third instar nymph, one fourth instar nymph, one fifth instar nymph, one adult male and one adult female. RNA was extracted using a Trizol protocol (Invitrogen).

### Genome sequencing and assembly

Genomic DNA was sequenced using HiSeq2500 Illumina technology. 180bp and 500bp paired-end and 3kb and 10kb mate-pair libraries were constructed and 100bp reads were sequenced. Estimated coverage was 28.6x, 7.3x, 21x, 17x, 72.9x respectively for each library. Sequenced reads were assembled in draft assembly using ALLPATHS-LG ^96^ and automatically annotated using custom MAKER2 annotation pipeline ^97^. (More details can be found in Supplementary Data). Expected genome size was calculated counting from Kmer based methods and using Jellyfish 2.2.3 and perl scripts from https://github.com/josephryan/estimate_genome_size.pl.

### Community curation of the *G. buenoi* genome

International groups within the i5k initiative have collaborated on manual curation of *G. buenoi* automatic annotation. These curators selected genes or gene families based on their own research interests and manually curated MAKER-predicted gene set GBUE_v0.5.3 at the i5k Workspace@NAL ^98^ resulting in the non-redundant Official Gene Set OGSv1.0 (Pending ADC URL).

### Assessing genome assembly and annotation completeness with BUSCOs

Genome assembly completeness was assessed using BUSCO ^28^. The Arthropoda gene set of 2 675 single copy genes was used to test *G. buenoi* predicted genes.

### Orthology analyses

OrthoDB8 (http://orthodb.org/) was used to find orthologues of *G. buenoi* (OGS 1.0) on 76 arthropod species. Proteins on each species were categorised using custom Perl scripts according to the number of hits on other eight arthropod species: *Drosophila melanogaster, Danaus plexippus, Tribolium castaneum, Apis mellifera, Acyrthosiphon pisum, Cimex lectularius, Pediculus humanus* and *Daphnia pulex*.

### Insulin receptors phylogeny

Sequences were retrieved from ‘nr’ database by sequence similarity using BLASTp with search restricted to Insecta (taxid:50557). Each *G. buenoi* InR sequence was individually blasted and best 250 hits were recovered. A total of 304 unique id sequences were retrieved and aligned with Clustal Omega ^99-101^ and a preliminary phylogeny was built using MrBayes ^102^ (one chain, 100 000 generations). Based on that preliminary phylogeny combined with Order and Family information, all isoforms could be confirmed and a representative(s) of each Order was selected (Supplementary Data File 3). Final InR phylogeny tree was estimated using MrBayes: four chains, for 500 000 generations and include InR sequences from *Acyrthosipon pisum* (2), *Aedes aegypti* (1), *Apis mellifera* (2), *Athalia rosae* (2), *Atta colombica* (2), *Bemisia tabici* (1), *Blatella germanica* (1), *Danaus plexippus* (1), *Diaphorina citri* (1), *Fopius arisanus* (2), *Halyomorpha halys* (2), *Nilaparvata lugens* (2), *Pediculus humanus* (1), *Tribolium castaneum* (2) and *Zootermopsis nevadensis* (2).

### Cytochrome P450 proteins phylogeny

CYPs phylogenetic analysis was performed using Maximum-Likelihood method and the trees were generated by MEGA 6. The phylogenetic trees were generated by MEGA 6 with Maximum-Likelihood method using the amino acid sequences from *Gerris buenoi* (Gb), *Rhodnius prolixus* (Rp), *Nilaparvata lugens* (Nl), Bombyx mori (Bm) *and Tribolium castaneum* (Tc). All nodes have significant bootstrap support based on 1000 replicates.

## Declarations

### Ethics approval and consent to participate

Not applicable

### Consent for publication

Not applicable

### Availability of data and material

GenBank assembly accession: GCA_001010745.1

### Competing interests

The authors declare that they have no competing interests

### Funding

ERC CoG grant #616346 to A.K., Genome sequencing, assembly and automated annotation was funded by a grant U54 HG003273 from the National Human Genome Research Institute (NHGRI to R.A.G.), German Research Foundation (DFG) grants PA 2044/1-1 and SFB 680 to K.A.P., NSERC Postdoctoral Fellowship grant to R.R.

## Authors’ contributions

D.A., A.K., and S.R. conceived, managed, and coordinated the project; S.D., D.M.M., R.A.G. and S.R. managed the i5k project; S.L.L. led the submissions team; HV.D., H.C. led the library team; Y.H., H.D. led sequencing team; J.Q., S.C.M., D.S.T.H., S.R. and K.C.W. assembled the genome; D.S.T.H. and S.R. did the automated annotation; D.A., R.R., M.F., J.B.B., H.M.R., K.A.P., S-J.A., M.M-T., E.A., F.B., T.C., C.P.C., A.G.C., A.J.J.C., A.D., E.M.D., E.D., E.N.E., M-J.F., C.G.C.J., A.J., E.J.J., J.W.J., M.P.L., M.L., A.M., B.O., A.L-O., A.R., P.N.R., A.J.R., M.E.S., W.T., M.v.d.Z., I.M.V.J., A.V.L. and S.V. contributed to manual curation; LM.C., C.P.C., M.M.T and M.F.P. performed curation quality control and generated the OGS; E.A., M-J.F. contributed to statistical analyses of wing development and polyphenism genes; I.M.V.J. and K.A.P. performed associated phylogenetic and synteny analyses of Wnt and Homeodomain transcription factor gene clusters; D.A. performed InR phylogeny and InR analyses; R.R. contributed with Insulin/TOR signaling genes analyses; M.F. perfomed opsins analyses; J.B.B. analyzed aquaporins, cuticle genes and antioxidant proteins; H.M.R. carried out chemoreceptors analyses; K.A.P. and I.M.V.J. perfomed Homeobox genes and Wnt signaling pathway analyses; S-J.A. analyzed UGTs detoxification genes; E.A. conducted wing poliphenism analyses; F.B. performed nuclear receptors and bHLH-PAS proteins analyses; A.G.C. carried out early developmental genes analyses; E.D. conducted histone genes and histone modification machinery; E.N.E., A.M. and B.O. performed cysteine peptidase analyses; C.F. contributed to analysis of Beadex duplication; A.J. analyzed Homeobox transcription factors; C.G.C.J. and M.v.d.Z. performed immune genes analyses; R.R., M.F., J.B.B., H.M.R., K.A.P., S-J.A., E.A., F.B., A.G.C., E.D., E.N.E., C.F., A.J., B.O., A.M., A.L-O., M.v.d.Z. and I.M.V.J. contributed to report writing; D.A. and A.K. wrote the manuscript; D.A., A.K., F.B., M.F. J.B.B., H.M.R, K.A.P. and S.R. edited the manuscript; D.A. and A.K. organized the Supplementary Materials. All authors approved the final manuscript

## Acknowledgements

We thank the staff at the Baylor College of Medicine Human Genome Sequencing Center for their contributions. Mention of trade names or commercial products in this publication is solely for the purpose of providing specific information and does not imply recommendation or endorsement by the U.S. Department of Agriculture. USDA is an equal opportunity provider and employer. We thank Jack Scanlan for annotating Ecdysteroid kinase family. We thank Lois Taulelle and Hervé Gilquin for providing access to computing resources in the Pôle Scientifique de Modélisation Numérique (PSMN) at the ENS Lyon. We thank Daniel Sloan for providing access to later version of *Pachypsylla venusta* opsin repertoire in addition to earlier genome assembly ^103^.

